# Gene-level metagenomics identifies genome islands associated with immunotherapy response

**DOI:** 10.1101/2020.10.09.333971

**Authors:** Samuel S. Minot, Kevin C. Barry, Caroline Kasman, Jonathan L. Golob, Amy D. Willis

## Abstract

Researchers must be able to generate experimentally testable hypotheses from sequencing-based observational microbiome experiments to discover the mechanisms underlying the influence of gut microbes on human health. We describe a novel bioinformatics tool for identifying testable hypotheses based on gene-level metagenomic analysis of WGS microbiome data (*geneshot*). By applying *geneshot* to two independent previously published cohorts, we identified microbial genomic islands consistently associated with response to immune checkpoint inhibitor (ICI)-based cancer treatment in culturable type strains. The identified genomic islands are within operons involved in type II secretion, TonB-dependent transport, and bacteriophage growth. These results, as well as the underlying methodology, inform further mechanistic studies and facilitate the development of microbiome-enhanced therapeutics.

---

Observational studies on the human gut microbiome using next-generation sequencing (whether 16S rRNA amplicons or whole genomic sequencing) have firmly established that the human gut microbiome affects health, disease, and response to treatments. The current analysis methods for whole-genome shotgun (or “WGS”) microbiome data have struggled to identify the underlying causal mechanisms, which are the critical next step to translate microbiome science into novel therapies. Identifying candidate mechanisms from observational studies is challenging because of the immense diversity of bacteria^1^, the limitations of reference genome databases^2^, widespread horizontal gene transfer^3^, and the difficulty of accurately predicting the biological function of microbial genes^4^.

One proposed solution is gene-level metagenomics: analysis with respect to abundance and presence of microbial protein-coding genes ^5^, but this approach has been limited by the high dimensionality of the data generated by this tactic. There are millions of detectable microbial protein-coding genes in an experiment, leading to challenges both with computation burden and statistical analysis. Our solution to the high-dimensionality of microbial protein-coding genes is to identify groups of genes with correlated levels of relative abundance across specimens (Co-Abundant Gene groups, or CAGs)^6^. CAGs may correspond to the core genomes of species, combinations of co-abundant species, or accessory genomic elements such as transposons, bacteriophage, or so-called genome “islands” which vary across strains^7^.

We have incorporated the tactic of identifying and combining protein-coding microbial genes into CAGs into a computational analysis pipeline for gene-level metagenomic analysis called *geneshot. Geneshot* is an end-to-end analysis workflow for microbiome experiments using CAGs as the fundamental unit of analysis. In brief, WGS data from each specimen is preprocessed to remove human reads and assembled *de novo*; predicted protein-coding genes are then deduplicated to make a reference gene catalog; the abundance of each gene in each specimen is estimated by assignment of raw reads against that gene catalog after rigorous handling of multiply-mapping reads^8^; genes are grouped into CAGs based on their co-abundance observed across the samples; we attempt to identify the possible taxonomic origins (by alignment against RefSeq) and functionally (using eggNOG mapper) of each gene; and the association of each CAG with experimental or clinical covariates is estimated (Fig. 1A). We consider if the CAGs associated with an outcome have member genes commonly represented by specific taxa. We used these identified taxa to identify operons associated with the outcome of interest, by aligning the member genes of associated CAGs to the reference genomes for that taxon.

**Figure 1.**
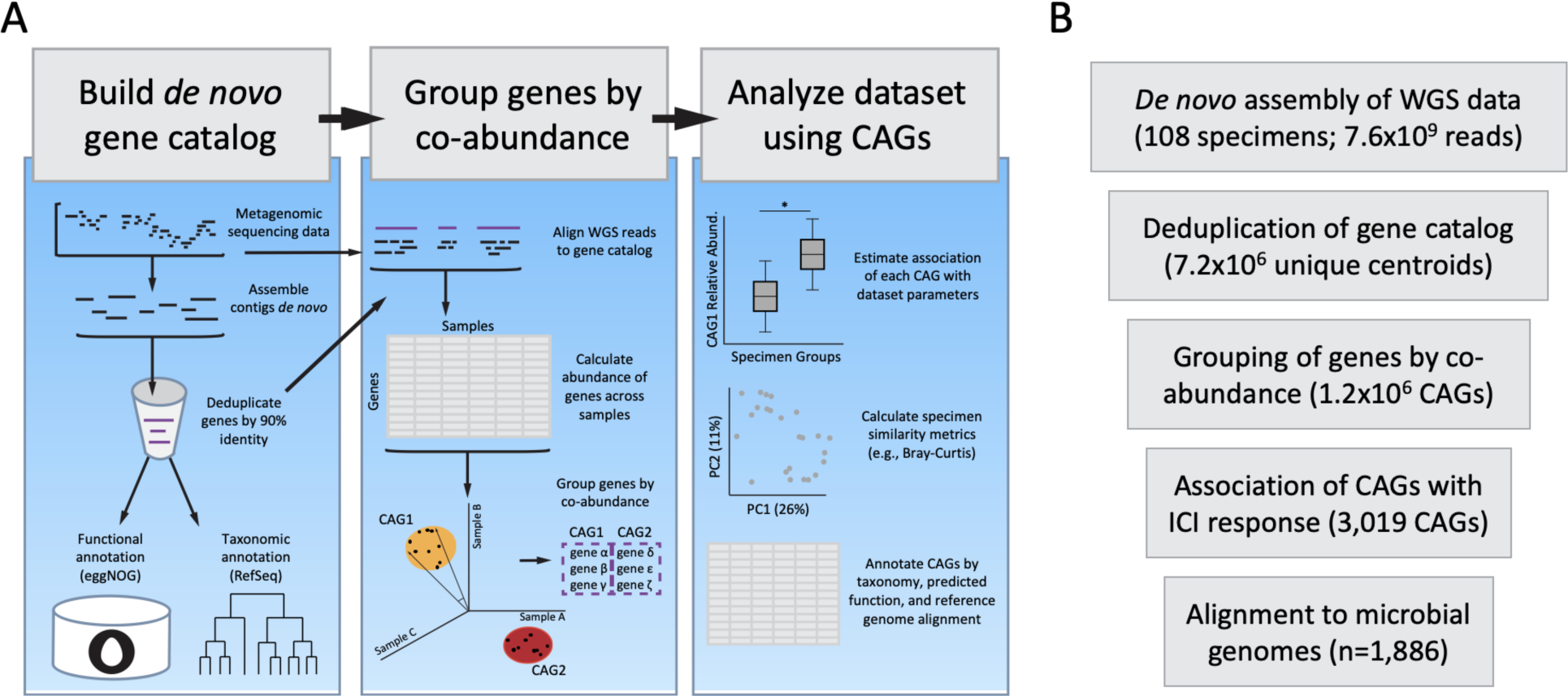
Schematic diagram of the *geneshot* tool for gene-level metagenomic analysis of microbiome experiments. A) Description of the data flow through *geneshot*, with the coordinated execution of multiple constituent bioinformatic processes in order to analyze experimental data on the basis of protein-coding genes encoded by microbes present in each physical specimen. B) Summary of the *geneshot* analysis to identify microbes associated with ICI response from published WGS datasets, indicating the number of biological features identified at each step.

In this study we employ *geneshot* to reanalyze data from metagenomic studies in people who have received immune checkpoint inhibitors for cancer. Immune checkpoint inhibitors (ICIs) are a class of immunotherapies that can induce robust and long-lasting protection from cancer^9–11^. However, across cancer indications the majority of patients have no objective response to ICI treatment^9–11^. Identifying the mechanisms that regulate response to ICIs is an area of intense research, and multiple studies have demonstrated that the gut microbiome can regulate the anti-tumor response induced by ICI in distant tissues, such as the lung and skin^12–14^. Further, transfer of the gut microbiome from patients to gnotobiotic mice transfers ICI responsiveness to the new host^12,13,15^. Taxonomic analysis of these datasets to identify the microbes responsible for transferring ICI responsiveness has yielded inconsistent results across cohorts. It is possible that the microbes driving the ICI-response phenotype may truly vary across these studies (e.g., due to biogeographical variation in human microbiome composition or the methodologies used to classify responder status), but the identification of common microbial genomic features associated with ICI response across multiple cohorts using gene-level analysis will accelerate understanding the mechanistic basis of the transfer of ICI responsiveness by the microbiome.

We used *geneshot* to analyze published datasets from microbiome studies investigating the stool microbiome of participants receiving ICI therapy for metastatic melanoma treatment^12–14^. Our goal was to identify microbial gene groups (CAGs) which differed in relative abundance in ICI responders compared to non-responders (termed ‘progressors’), while allowing for differences in the baseline relative abundance of each CAG in the two cohorts. Each stool microbiome sample yielded 75,488 - 536,005 (median 280,065) microbial genes by *de novo* assembly which were deduplicated to form a gene catalog of 7,209,758 unique protein-coding sequences (Fig. 1B, Fig. S1A). The majority of raw WGS reads (median 86.1%) were uniquely assigned to that gene catalog by translated nucleotide alignment, and 380,202 - 2,071,835 (median 1,174,474) genes were detected in each sample (Fig. S1B-C). Co-abundance clustering of the genes yielded 1,232,769 distinct CAGs; member genes within the most abundant CAGs were commonly represented in the reference genomes of common gut residents (Fig. S1D-E, S2). With an FDR threshold of 0.01, 3,019 CAGs (4,509 genes in total) were found to be significantly associated with ICI response, with 2,634 CAGs associated with progression and 385 CAGs associated with response (Fig. S3). These associated CAGs had member genes commonly present in *Odoribacter splanchnicus* and *Gemmiger formicilis* (which were found to be more abundant in responders), as well as *Coprococcus comes* (which was more abundant in progressors) (Figs. S4, S5).

To provide genomic and functional context for these ICI-associated CAGs, we next aligned the dataset to 1,886 reference genomes from the taxonomic families shown in Figure S4C. Rather than being spread throughout the genome, the genes comprising the CAGs were found in contiguous genomic regions (“islands”) ranging in size from 10-35kb in size. ICI-associated CAGs aligning to *Odoribacter splanchnicus* DSM 220712 corresponded to strain-specific genomic islands encoding type II secretion and TonB-dependent transport, which were found at higher abundance in the gut of individuals who responded to ICI therapy across both cohorts (Fig. 2). Genomic islands in *Clostridium sp. HMb25* and *Faecalibacterium prausnitzii* were annotated as integrated prophages, with an additional genomic island in *F. prausnitzii* annotated with a CRISPR defense system, supporting a role of bacteriophage growth in the ICI-microbiome interaction (Fig. S6). Thus, *geneshot* was able to identify not only potentially relevant taxa, but also strain specific genomic islands. Furthermore, *geneshot* identified specific possible mechanisms (type II secretion, TonB-dependent transport, and phage) that are suitable for targeted hypothesis testing (e.g., in model systems).

**Figure 2.**
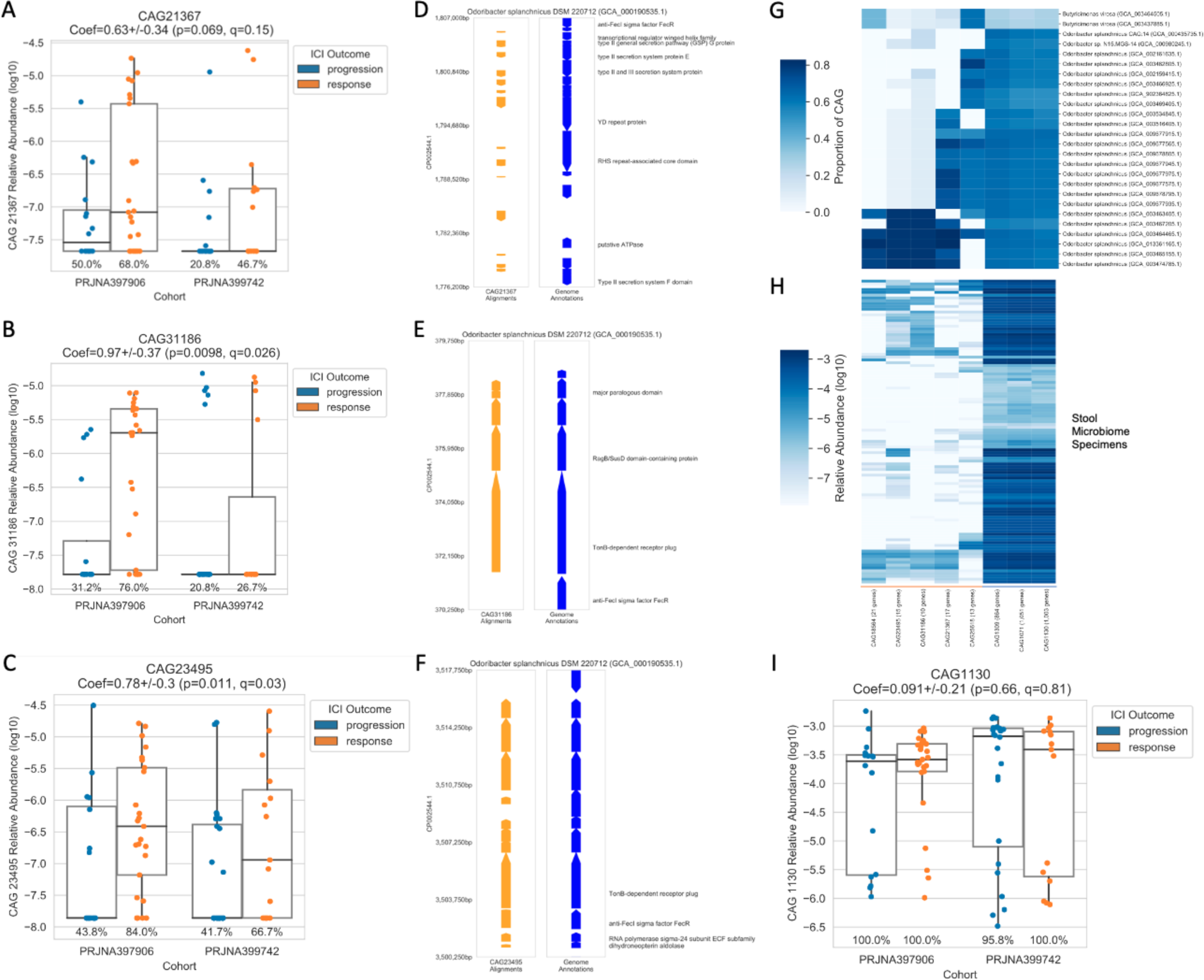
Genomic coordinates of ICI response-associated CAGs for *Odoribacter splanchnicus* DSM 220712. The relative abundance of each CAG (A, B, C, I) is shown across specimens, grouping specimens by cohort (x-position) and ICI response outcome (color). The genomic coordinates of individual CAGs (D, E, F) are shown with orange arrows for the region of alignment for each gene and blue arrows with text indicating the NCBI annotations in that region. The occurrence of CAGs across multiple reference genomes (G) as well as stool microbiome specimens from this dataset (H) differentiates the core genomic elements (underlined in blue) from the accessory (or strain-specific) genetic elements (underlined in orange).

In contrast to other end-to-end pipelines, the unit of analysis that underlies *geneshot* is *de novo* assembled protein-coding gene sequences, which are dimension-reduced via co-abundance clustering. One advantage of this gene-level approach to metagenomic analysis is a reduced reliance on reference databases (which are often incomplete) or inaccurate ontological hierarchies (e.g. the incompatibility of taxonomy with homoplasmy or horizontally transferred genetic elements). Moreover, by implementing an improved method for CAG construction *geneshot* is able to analyze gene-level abundances without being restricted to organismal groupings (such as MAGs or “metagenomic species”).

*Geneshot’s* integration of reference and taxonomic databases is oriented around deriving maximum benefit from the sparsely sequenced and annotated uncultivated organisms within the gut. After identifying CAGs associated with an outcome, we then consider where the member genes of the associated CAGs have been observed in reference genomes. The actual organism representing these genes in a given specimen need not have a reference genome, just a somewhat related genomic island. Thus, *geneshot* maximizes what can be inferred from reference databases without being dependent upon the same references for identification of relevant genomic islands.

Finally, by wrapping together the set of tasks required for comprehensive gene-level metagenomic analysis using the Nextflow workflow management system^16^ we provide a convenient mechanism for microbiome analysis which can be implemented across a variety of high-performance computing infrastructures with minimal configuration required for each user. We hope that *geneshot* can be used widely to enhance the quality of microbiome research, while also providing insights which may contribute to the development of microbiome-based therapeutics for cancer and other human diseases.

## Methods

### 1. Bioinformatic approach of *geneshot* for gene-level metagenomics

The process of performing gene-level metagenomic analysis is implemented by *geneshot* using a series of bioinformatic processes which are all coordinated into an overarching workflow using the free and open source Nextflow^16^ workflow management system. By providing options as “flags” to the workflow users can modify the details of how the analysis is executed, and a record of the parameters used to execute the workflow can be saved (along with a summary of the computational resources which were used) with the reporting functionality provided by Nextflow. The complete set of code which constitutes *geneshot* can be found in the GitHub repository https://github.com/Golob-Minot/geneshot under the open source MIT License, with documentation provided in the associated wiki.

The overview of the analytical tasks performed by *geneshot* are as follows (see section 6 for a complete reference of the component tools referenced here):

1. Input WGS reads are preprocessed with adapter trimming using barcodecop and cutadapt and host reads are removed by subtractive alignment using BWA. All of the following steps use the FASTQ files which are output by this preprocessing step.
2. WGS reads from each specimen are *de novo* assembled individually using MegaHit and coding regions are predicted using Prodigal.
3. The conceptual translations of every predicted coding region are deduplicated across all specimens using a combination of linclust and DIAMOND, each of which applies a fixed threshold of amino acid similarity (default: 90%) and alignment coverage (default: 50%) to retain only the centroids from each cluster of coding sequences.
4. The preprocessed WGS reads from step 1 are then aligned against the gene catalog from step 3 in order to estimate the relative abundance of the organisms in each specimen which encode each gene. The alignment is carried out with DIAMOND (in blastx mode). Those alignments are then processed by FAMLI in order to resolve any reads which align equally well to multiple references by picking a single reference using an expectation maximization approach that uses a target metric of evenness of sequencing coverage across each reference as the target metric^8^.
5. The relative abundance of each gene from the catalog across each specimen is computed as the depth of sequencing across each gene (the number of bases from WGS reads aligned to each gene divided by the length of the gene) divided by the sum of sequencing depth across all genes for that specimen. This provides a relative abundance estimate for the genomes encoding each gene in the source specimen that is adjusted for gene length.
6. Using the relative abundance of each gene across each specimen, genes are clustered using average linkage clustering and the cosine distance metric, applying a fixed distance metric to join genes into Co-Abundant Gene Groups (CAGs). Each gene from the catalog which is detected in at least one sample belongs to one and only one CAG. Two complementary approaches are used to provide a computationally tractable approach to this process: (a) genes are clustered in sub-groups (shards) which are iteratively combined over five sequential rounds; and (b) an Approximate Nearest Neighbor algorithm is used to rapidly index and retrieve co-abundant gene ‘neighborhoods’ to limit the search space for expensive computation of pairwise distance matrices as described previously^6^. To mitigate any risk of the use of shards in (a) which could find sub-optimal clusters in early rounds, the stringency of the cosine distance threshold used to identify clusters is increased by a factor of 2 in all pre-clustering steps. Only in the final round of clustering is the user-specified cosine distance threshold applied, which results in the formation of CAGs which are the superset of the stringent groups formed in earlier rounds.
7. Independently from steps 4-6, the gene catalog may be optionally annotated by taxonomic classification using DIAMOND (which assigns the lowest common ancestor of top hits in blastp mode), as well as functional annotation using eggNOG-mapper. These annotation steps are optional and are only executed if the user provides the reference databases required by either tool.
8. If the user provides a formula for statistical analysis using the variables defined in the manifest (the input file which is also used to label each pair of WGS FASTQ files with the appropriate specimen name), then the number of reads mapped to each CAG will be modeled using a beta-binomial model with that formula describing the logit of the expected relative abundance of the CAG. The model is fit using corncob^17^. If taxonomic annotation was performed in step 7, taxon abundance coefficients are determined by aggregating the corncob results over all CAGs for which any constituent gene is taxonomically annotated as that taxon. The taxon-level estimated coefficients are then modelled using the errors-in-response model betta^18^ with no additional covariates (i.e., an intercept-only model is fit), and the hypothesis that the intercept is zero is tested. This approach allows an overall assessment of the effect of the covariate on the abundance of each taxon in a CAG-based analysis. Similarly, if functional annotation was performed in step 7, the same approach using betta and corncob is applied to all CAGs sharing the same functional annotation, which allows an overall assessment of the effect of the covariate on the abundance of each function. Taxon abundance coefficients are used in this analysis to prioritize reference genomes for more detailed analysis by exhaustive alignment of the genes from each CAG against individual reference genomes.
9. In the final step all of the outputs from each step are aggregated into HDF5 format, which organizes multiple tables into a single file using an internal organization structure to identify each of the elements of the results. The updated description of all data output by *geneshot* can be found in the associated documentation (https://github.com/Golob-Minot/geneshot/wiki/Output-Files).

### 2. Preparing inputs for *geneshot*

In order to run *geneshot*, users must prepare (1) a manifest in CSV format with columns for “specimen”, “R1”, “R2”, and any other additional metadata needed for statistical analysis, and (2) the set of FASTQ files (gzip-compressed) listed in the manifest, with forward and reverse reads listed separately in the “R1” and “R2” columns, respectively. In this format, a single specimen may be made up of multiple FASTQ file pairs by listing those files across multiple rows.

### 3. Setting up Nextflow

In order to run *geneshot* on the available computational resources, start by installing the workflow management tool Nextflow and configuring it. The complete documentation for Nextflow can be found at https://www.nextflow.io/docs/latest/getstarted.html. The process of configuring Nextflow is accomplished by saving a “nextflow.config” file in the user’s home directory which describes the compute resources which should be used (local execution, SLURM, PBS, AWS, etc.), and which will be referenced when running *geneshot*.

### 4. Running *geneshot*

The complete set of instructions for running *geneshot* can be found on the wiki associated with the tool at https://github.com/Golob-Minot/geneshot/wiki. In order to run *geneshot*, the user starts with the command “nextflow run Golob-Minot/geneshot” and appends any of the flags shown in Supplementary Table 1 (e.g. --manifest manifest_file.csv).

### 5. Reference databases

Annotation of the gene catalog generated by *geneshot* is performed using reference databases for DIAMOND and eggNOG-mapper. The databases used for this analysis were generated based on the documentation in the associated tools, and are available on AWS S3 at the following locations:

- Taxonomic annotation is based on a DIAMOND index of NCBI’s RefSeq genome database generated on January 15, 2020: s3://fh-ctr-public-reference-data/tool_specific_data/geneshot/2020-01-15-geneshot/DB.refseq.tax.dmnd
- Functional annotation is based on eggNOG-mapper v5.0 database downloaded on July 17th, 2020: s3://fh-ctr-public-reference-data/tool_specific_data/geneshot/2020-06-17-eggNOG-v5.0/eggnog_proteins.dmnd and s3://fh-ctr-public-reference-data/tool_specific_data/geneshot/2020-06-17-eggNOG-v5.0/eggnog.db

### 6. Bioinformatic component tools and dependencies

All of the component tools used by *geneshot* are available publicly as Docker images which are automatically generated from publicly available Dockerfile source code. The images used by the current version of *geneshot* are listed in Supplementary Table 2.

### 8. Genome alignments

The process of aligning genes from the *geneshot* output against a set of reference genomes was executed the Annotation of Microbial Genomes by Microbiome Association (AMGMA) tool. This tool aligns genes from the *geneshot* gene catalog against a user-specified set of microbial genomes and then calculates summary metrics for each genome including the proportion of the genome which is aligned by genes which are associated with a given parameter. The genome alignment figures in this manuscript show the position of the genome alignment for the set of genes contained in the CAG of interest, as well as the location and annotation of genes recorded in GFF format within the NCBI RefSeq database.

Alignment is performed using DIAMOND (using the dependencies shown in Table S2) with the complete set of Python code used to aggregate alignment results available at https://github.com/FredHutch/AMGMA/. An example of alignment to *de novo* assembled contigs, performed with the same set of steps, is shown in Fig. S7.

### 9. Datasets used for analysis

The data analyzed in this manuscript consists of the publicly available FASTQ records and associated metadata available for the BioProjects PRJEB22893^12^ (n=25), PRJNA397906^14^ (n=44), and PRJNA399742^13^ (n=172). The public metadata for PRJEB22893^12^ did not include any annotation of ICI response phenotype, and so was omitted from the statistical model used to analyze CAG abundances. However, the CAG abundance data for that set of samples is present in the raw results from the *geneshot* results published alongside this paper.

### 10. Data availability

The complete set of *geneshot* results described in this manuscript are available via FigShare (accession forthcoming) in the format described in https://github.com/Golob-Minot/geneshot/wiki/Output-Files. Genome alignments generated by the AMGMA tool described in this manuscript are also available in that repository in self-documenting HDF5 format.

## Acknowledgements

We would like to acknowledge our funding support including NIGMS R35 GM133420-01 (PI: Dr. Willis) and the Fred Hutch Microbiome Research Initiative.

## Author Contributions

SM, JG, and AW contributed to the conceptual approach of the *geneshot* software. SM primarily executed the development of the *geneshot* software, incorporating code contributed by JG and AW. SM performed the computational analysis of public ICI datasets. All authors (SM, KB, CK, JG, and AW) contributed to the interpretation of results, development of figures, and drafting of manuscript text.

## Supplementary Figures

**Figure S1.**
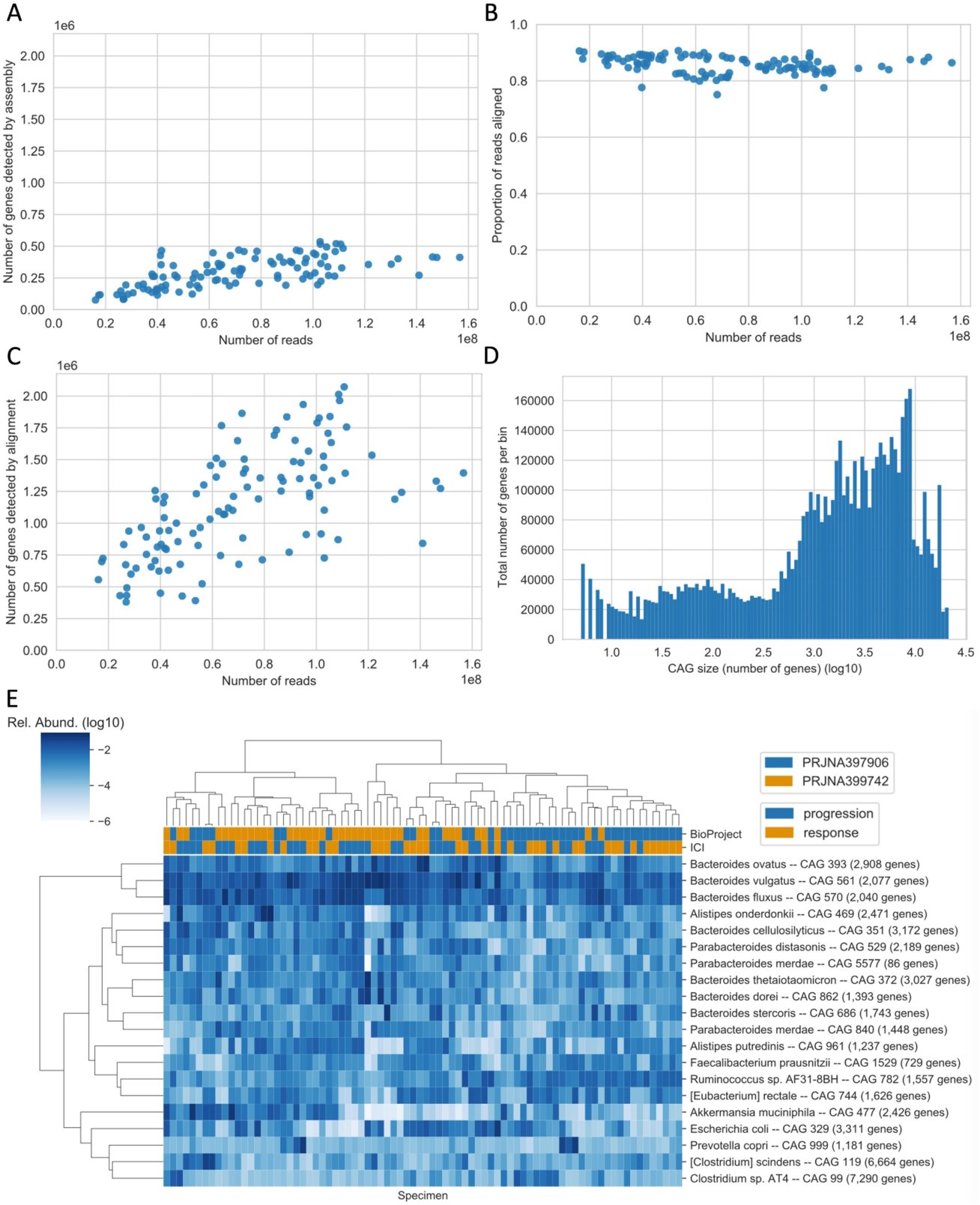
Summary of gene-level metagenomic analysis results generated by *geneshot*. The results for each specimen across all cohorts is shown in comparison to the number of sequence reads for each specimen (x-position) including the number of genes detected by *de novo* assembly of each individual specimen (A), the proportion of reads from each specimen which align uniquely to the *de novo* gene catalog (B), and the number of genes from the gene catalog detected by alignment (C). The distribution of genes across CAGs of different sizes is shown in (D), with the horizontal axis showing the CAG size range and the vertical axis showing the number of genes belonging to CAGs within that size range. (E) shows the relative abundance (log10) of the CAGs with the highest relative abundance (rows, annotated by majority taxonomic annotation) across all specimens (columns, annotated by BioProject and ICI response label), with hierarchical clustering by relative abundance across rows and columns.

**Figure S2.**
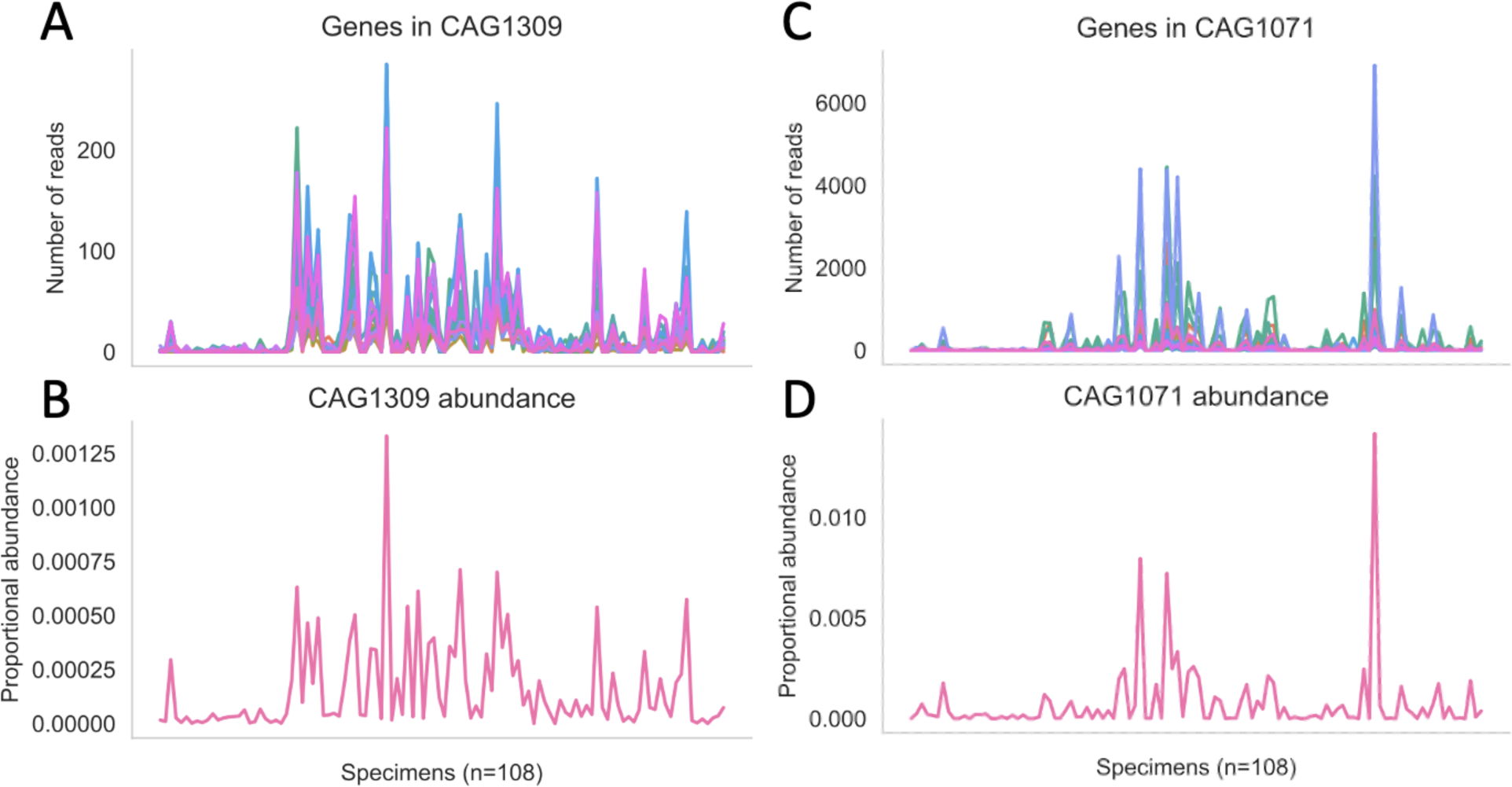
Visual depiction of CAG-level abundance calculation. Each CAG consists of a group of genes, each of which is detected in each specimen by alignment of WGS reads. The abundance of genes from each CAG are shown in (A) and (C) in units of the absolute number of reads aligned uniquely from each specimen. The ‘depth’ of sequencing of each gene is a commonly-used length-normalized measure in which the total number of bases from all aligned reads is divided by the length of each gene. Genes are grouped into CAGs based on average linkage clustering using the cosine distance metric computed from the vector of sequencing depth values across all specimens. The abundance of each CAG is computed as the sum of sequencing depth for all genes in the CAG, divided by the sum of sequencing depth of all genes detected. That proportional abundance metric is shown for each CAG in (B) and (D). Specimens are arranged in alphabetical order on the horizontal axis, with abundance measures for each specimen on the vertical axis.

**Figure S3.**
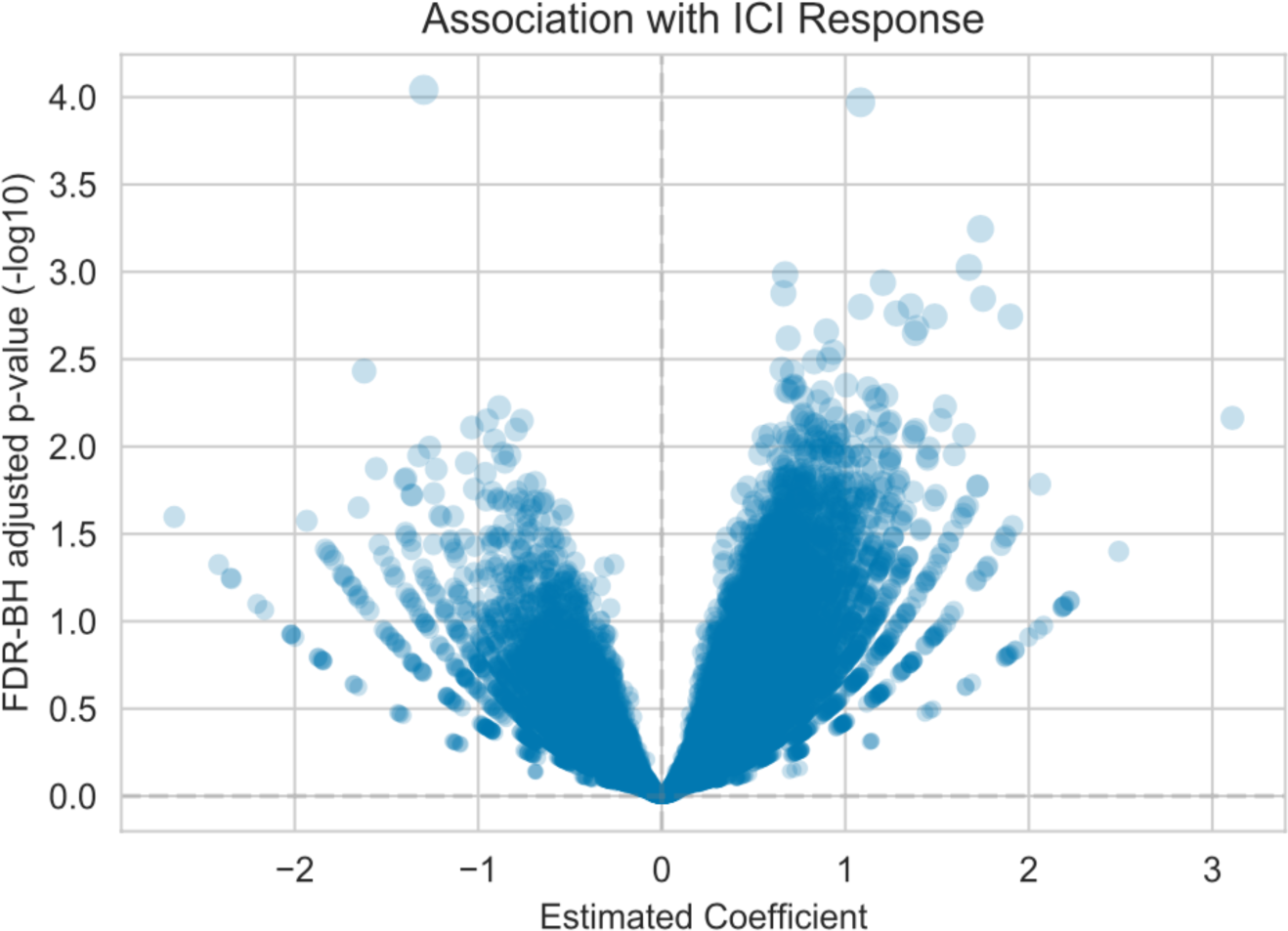
Volcano plot showing the distribution of association estimates for CAGs with ICI response. Each point represents a single CAG, showing all CAGs which contain at least five constituent genes. The horizontal axis shows the estimated coefficient of association with ICI response (positive values indicate a higher relative abundance in ICI responders in comparison to progressors). The vertical axis shows the *p*-value for that coefficient of association for each CAG after adjustment for multiple hypothesis testing with the FDR-BH procedure.

**Figure S4.**
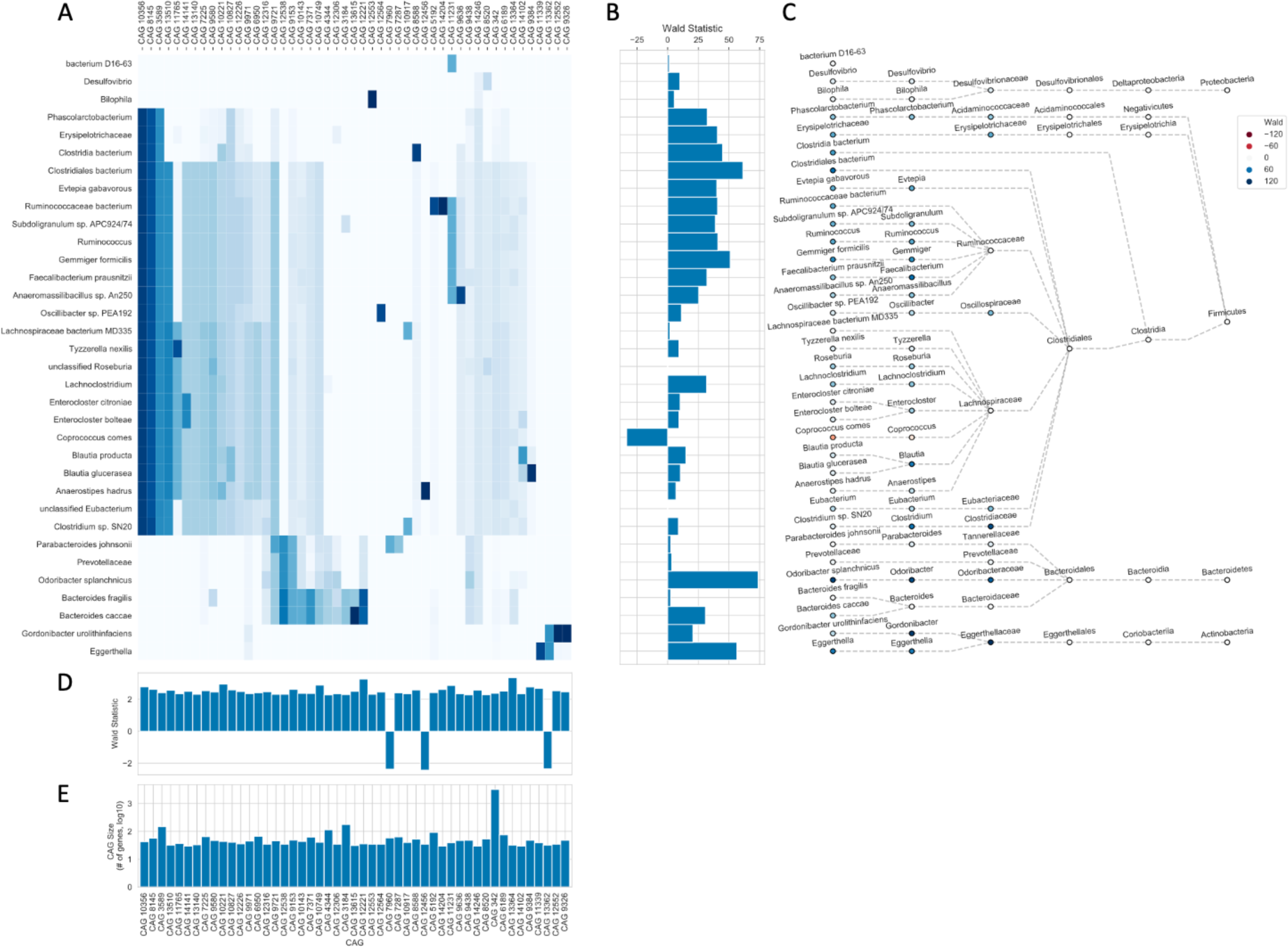
Taxonomic annotation of CAGs associated with ICI response. The taxonomic annotation of genes found in the 50 CAGs (columns) which were most consistently associated with ICI response (after applying a minimum size threshold of 30 genes), are shown in (A) with shading indicating the proportion of genes assigned to the indicated taxon (rows). The Wald statistic summarizing the association with ICI response is shown for taxa in (B) and for individual CAGs in (D). A taxonomic tree annotating those organisms is shown in (C), and CAG size is shown in (G).

**Figure S5.**
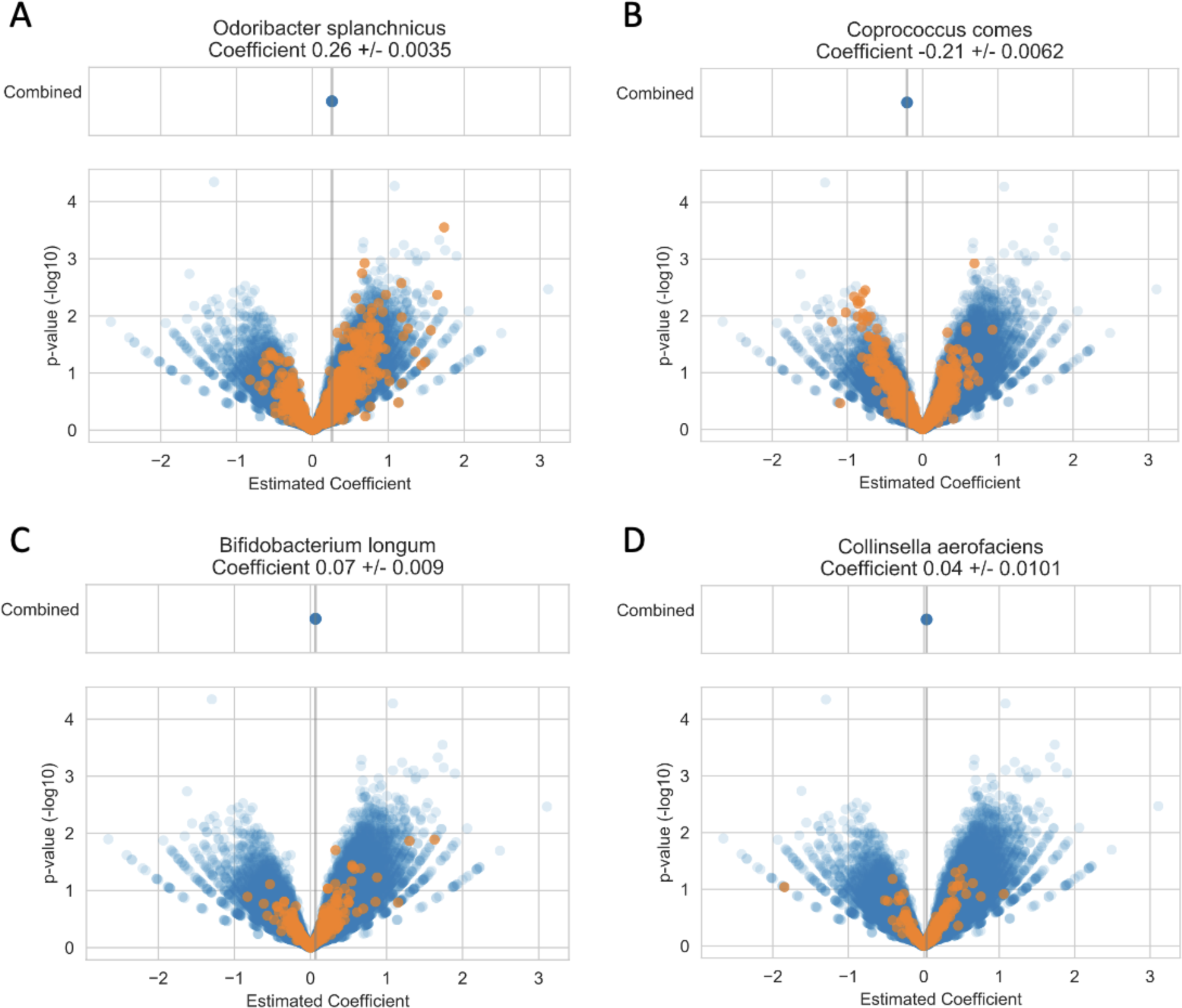
Visual depiction of the aggregation of CAG-level ICI-response association metrics by taxon. For each organism, the lower panel shows the estimated coefficient and -log10 *p*-value for the association of each CAG with ICI response. Every CAG either contains at least one gene which was taxonomically annotated to the indicated organism (orange), or there were no genes in that CAG with that taxonomic annotation (blue). The aggregate taxon-level ICI-response association for each organism is shown in the upper panel from each subplot, with the estimated coefficient +/- standard deviation indicated in the subplot title. The vertical grey line in each subplot marks the taxon-level estimated coefficient value.

**Figure S6.**
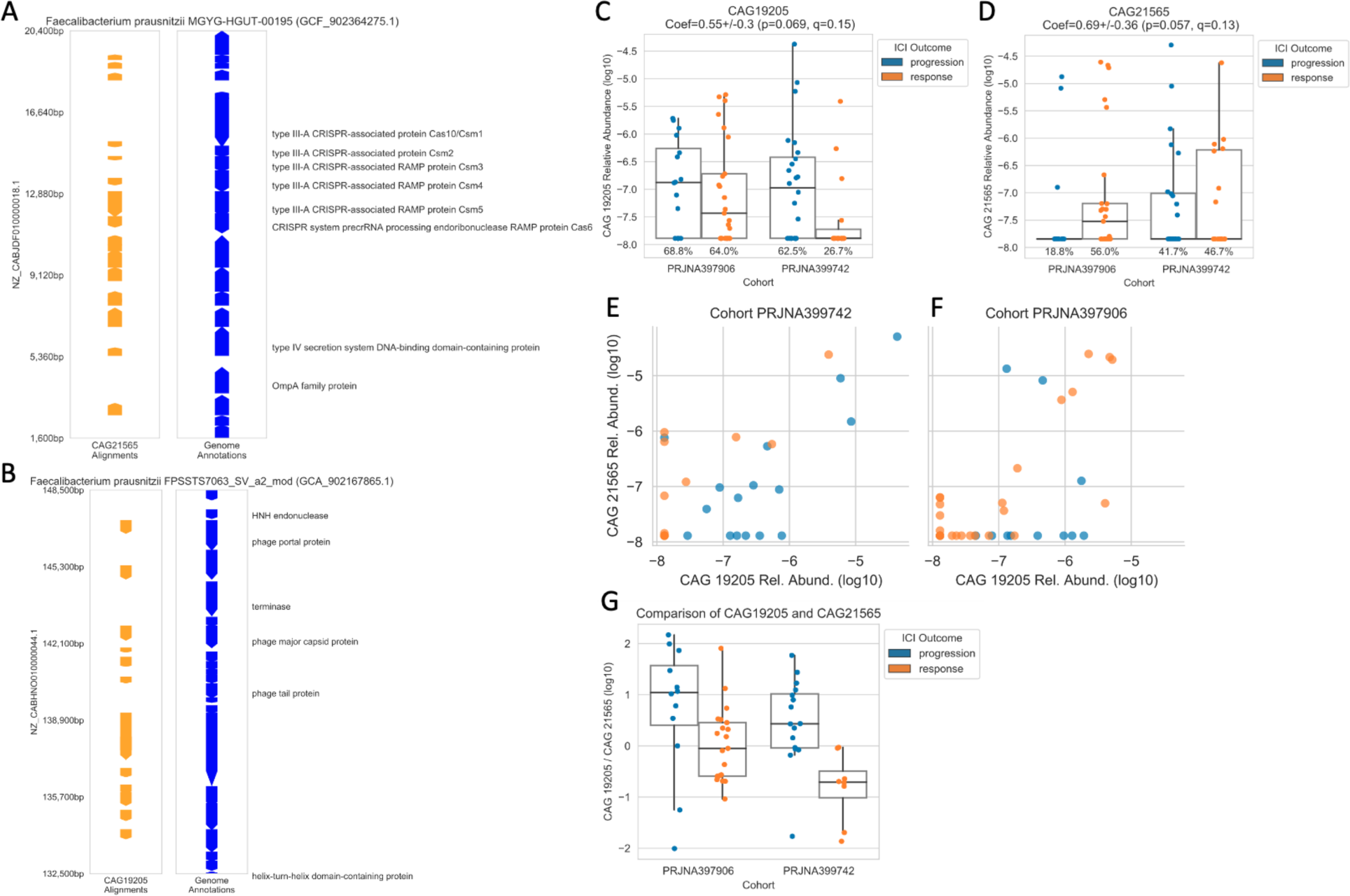
A putative bacteriophage of *Faecalibacterium prausnitzii* is associated with ICI non-response, in contrast to an associated bacteriophage defense island that is associated with ICI response. The genomic coordinates of ICI response-associated CAGs are shown for two strains of *F. prausnitzii* (as in Figure 2) (A-B). In addition, the relative abundance of these two CAGs is compared directly (C-F), and the ratio of relative abundances is shown as a function of ICI response across both cohorts (G).

**Figure S7.**
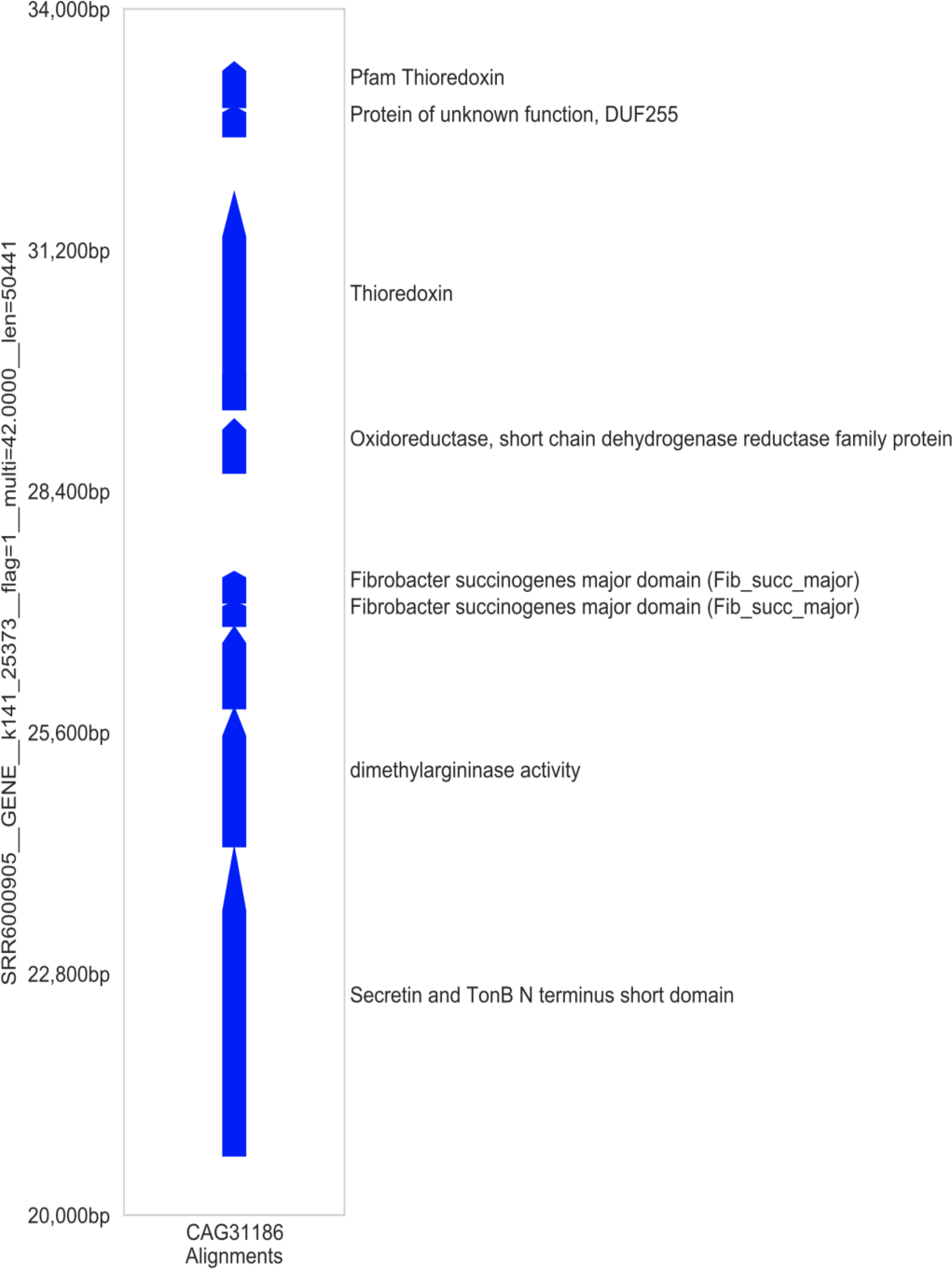
Example of CAG alignment to *de novo* assembly. The alignment coordinates for the alignment of a single CAG against the contigs generated by *de novo* assembly from this dataset is shown alongside the eggNOG functional annotations for the underlying genes. All alignments covered the entirety of the gene catalog sequence at >99% amino acid identity. The 14kb physical region shown in this plot encompasses all such high quality alignments for this CAG in this specimen.

**Table S1.**
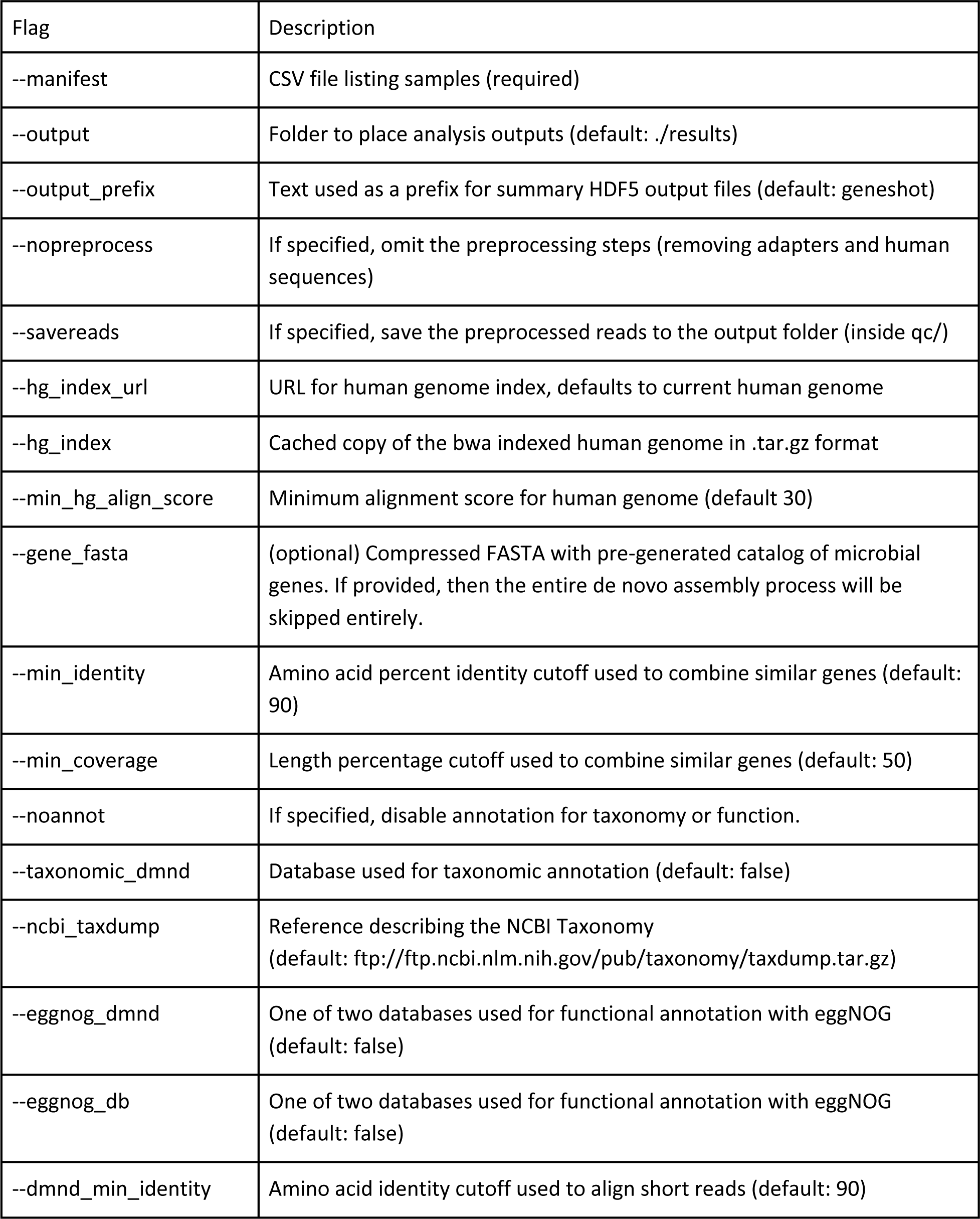
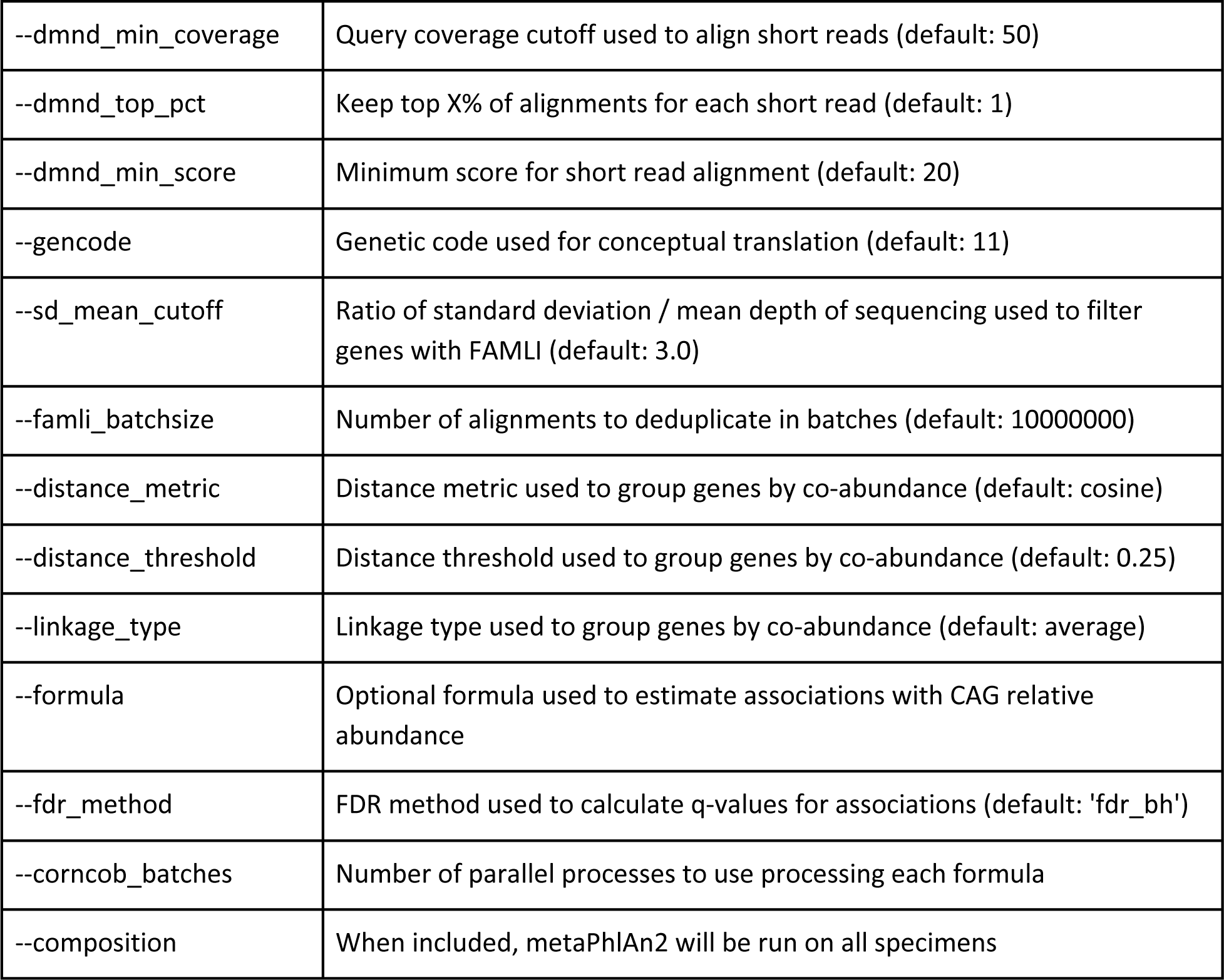
Description of flags used to customize parameters for running *geneshot*.

**Table S2.**
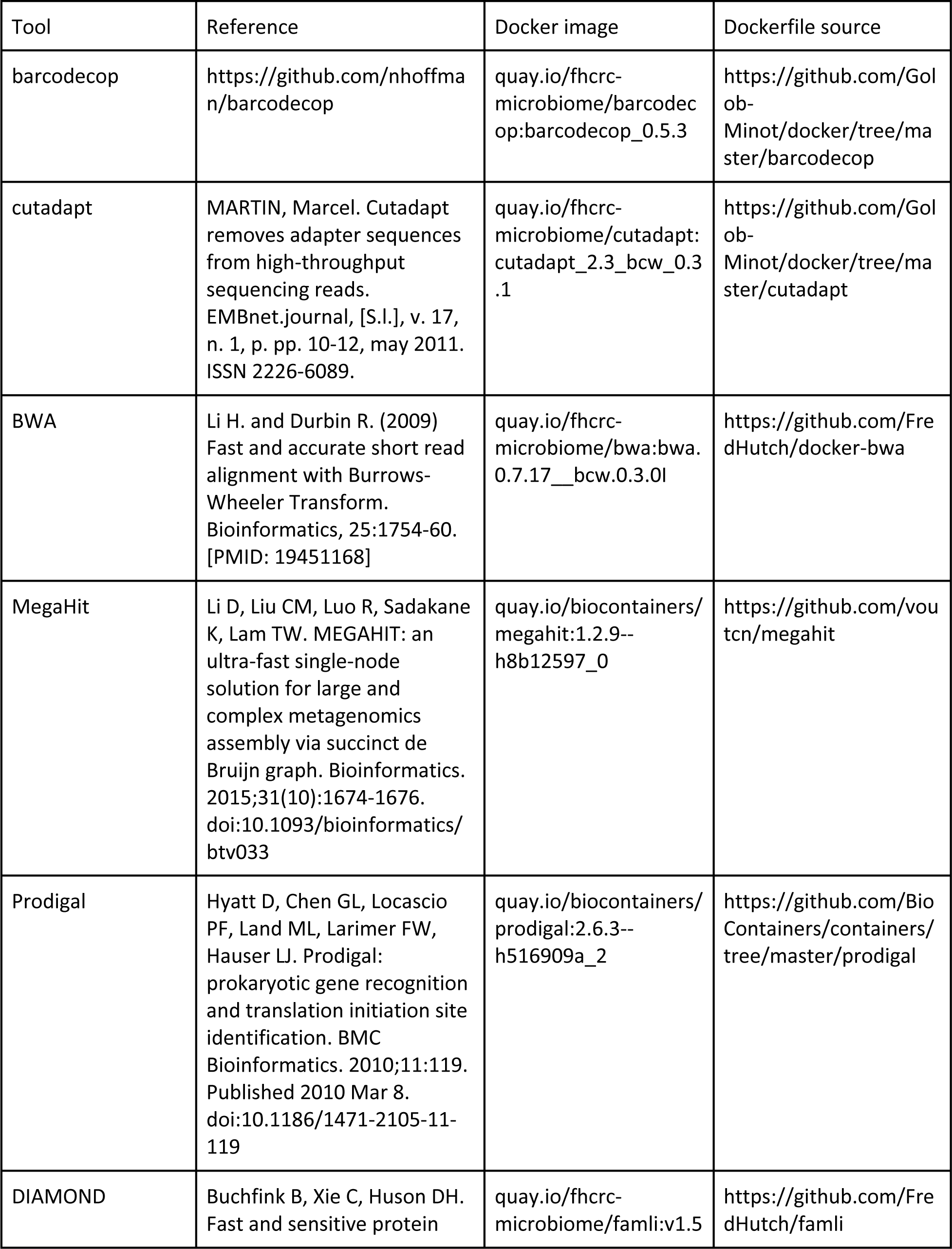
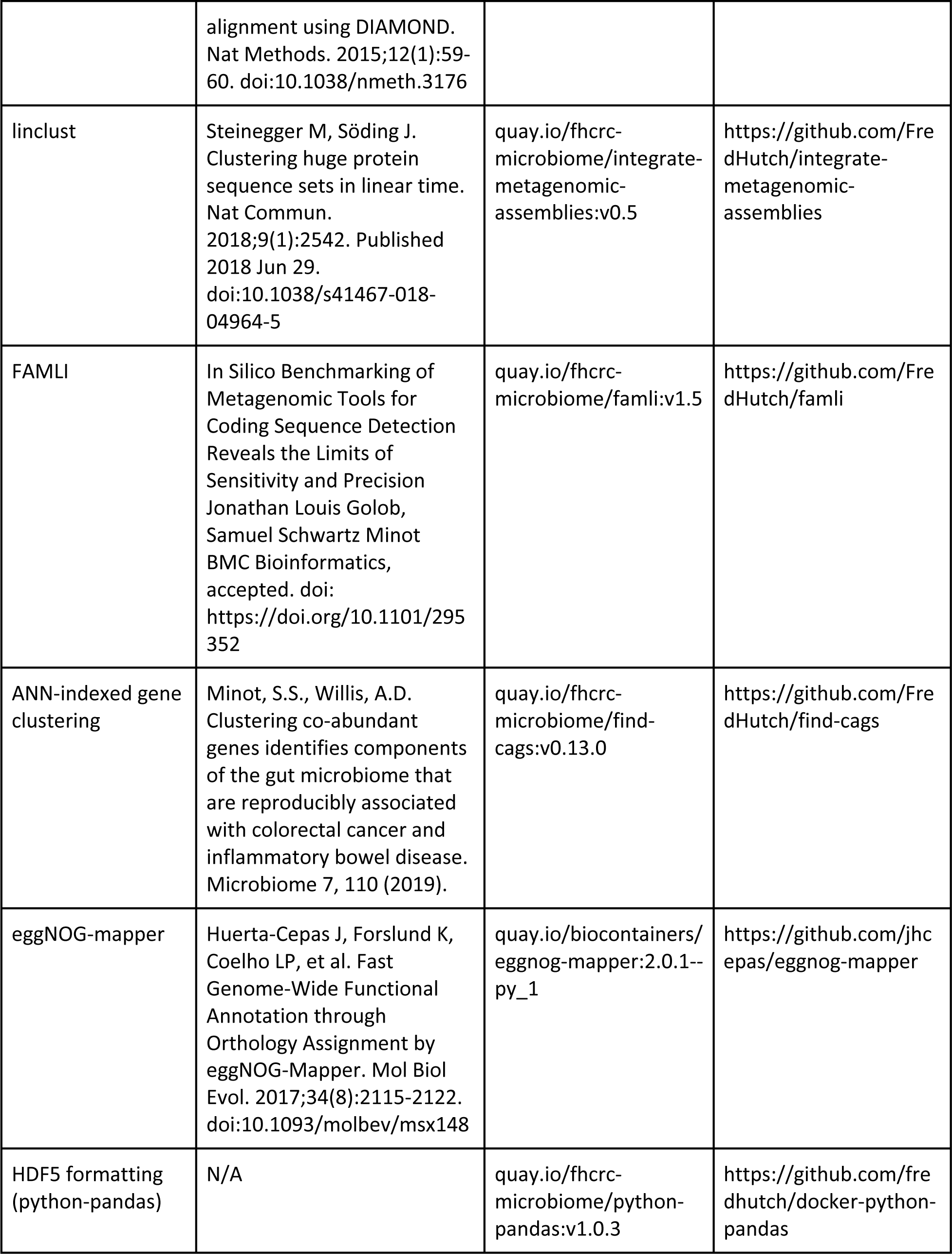
Description of software dependencies and versions used by *geneshot*.

